# The *Neurospora crassa* standard Oak Ridge background exhibits an atypically efficient meiotic silencing by unpaired DNA

**DOI:** 10.1101/333476

**Authors:** Dev Ashish Giri, Ajith V. Pankajam, Koodali T. Nishant, Durgadas P. Kasbekar

## Abstract

Meiotic silencing by unpaired DNA (MSUD) was discovered in crosses made in the standard Oak Ridge (OR) genetic background of *Neurospora crassa*. However, MSUD often was decidedly less efficient when the OR-derived MSUD tester strains were crossed with wild-isolated strains (W), which suggested either that sequence heterozygosity in *tester* x W crosses suppresses MSUD, or that OR represents the MSUD-conducive extreme in the range of genetic variation in MSUD efficiency. Our results support the latter model. MSUD was much less efficient in near-isogenic crosses made in a novel *N. crassa* B/S1 and the *N. tetrasperma* 85 genetic backgrounds. Possibly, regulatory cues that in other genetic backgrounds calibrate the MSUD response are missing from OR. The OR versus B/S1 difference appears to be determined by loci on chromosomes 1, 2, and 5. OR crosses heterozygous for a duplicated chromosome segment (*Dp*) have for long been known to exhibit an MSUD-dependent barren phenotype. However, inefficient MSUD in *N. tetrasperma* 85 made *Dp*-heterozygous crosses non-barren. This is germane to our earlier demonstration that *Dp*s can act as dominant suppressors of repeat-induced point mutation (RIP). Occasionally, during ascospore partitioning rare asci contained >8 nuclei, and round ascospores dispersed less efficiently than spindle-shaped ones.

**General abstract:** In crosses made in the standard OR genetic background of *Neurospora crassa*, an RNAi-mediated process called MSUD efficiently silences any gene not properly paired with its homologue during meiosis. We found that MSUD was not as efficient in comparable crosses made in the *N. crassa* B/S1 and *N. tetrasperma* 85 backgrounds, suggesting that efficient MSUD is not necessarily the norm in Neurospora. Indeed, using OR strains for genetic studies probably fortuitously facilitated the discovery of MSUD and its suppressors. As few as three unlinked loci appear to underlie the OR versus B/S1 difference in MSUD.

## Introduction

Meiotic silencing by unpaired DNA (MSUD) was discovered in crosses made in the standard Oak Ridge (OR) genetic background of *Neurospora crassa* (Aramayo and Metzenberg 1996; Shiu *et al*., 2001; Perkins, 2004). Any gene not properly paired with its homologous sequence during meiosis is transcribed into ‘aberrant RNA’ that is then processed into single-stranded “MSUD-associated small interfering RNA” (masiRNA), which directs a silencing complex to degrade complementary mRNA, thus silencing the unpaired gene and other genes homologous to it (Hammond, 2017). The ::*act, ::asm-1, ::Bml^r^, ::mei-3*, and ::*r*^*+*^ MSUD tester strains contain an additional copy of the *act* (*actin*), *asm-1^+^* (*ascospore maturation-1*), *Bml β–tubulin*), *mei-3*, or *r*^*+*^ (*round ascospores*) gene inserted at an ectopic location. The ectopic copy is unpaired in a *tester* x OR cross and induces the production of masiRNA which degrades its complementary mRNA, and the resulting deficit of actin, ASM-1, β-tubulin, MEI-3, or R protein manifests as striking ascus or ascospore phenotypes, whereas in a homozygous *tester A* x *tester a* cross the ectopic copy is paired, therefore there is no MSUD, and ascus and ascospore development is normal (Raju *et al*., 2007).

Surprisingly, MSUD was not as efficient when the OR-derived tester strains (*tester*^*OR*^) were crossed with several wild-isolated *N. crassa* strains (Ramakrishnan *et al*., 2011). Of 80 wild-isolated strains tested in crosses with the ::*Bml*^*r*^ and ::*mei-3* testers, only eight, designated as the “OR” type, showed silencing phenotypes comparable to the *tester*^*OR*^ x OR crosses. Crosses with four wild strains designated as the “Sad” type did not show MSUD, and the remaining 68 strains showed an intermediate phenotype, in that, the crosses silenced *bml* but not *mei-3*^+^, and they were designated the “Esm” type. One hypothesis (model 1) to explain these results posits that sequence heterozygosity between the *tester*^*OR*^ and wild genomes either overwhelms the MSUD machinery or causes asynapsis and self-silencing of one or more MSUD gene. This model is based on the fact that deletion alleles of several MSUD genes act as dominant suppressors of MSUD (Samarajeewa *et al*., 2017 and references therein), presumably because they cause their wild-type homologue to become unpaired, induce its autogenous silencing, and thus “silence the silencer” (Hammond, 2017). The *sad-1Δ* and *sad-2Δ* deletions (i.e. *Sad-1* and *Sad-2* (*Suppressor of ascus dominance-1* and-*2*) were strong dominant suppressors whereas the other MSUD gene deletions were less effective, possibly because of high expression or long protein half-life (Hammond *et al*., 2013a; Decker *et al*., 2017). The *Sad-1* and *Sad-2* dominant MSUD suppressors suppressed the classical ascus-dominant mutants *Ban* (*Banana*), *Dip-1* (*Diploid ascospores-1*), *Pk*^D^ (*Peak-Dominant*), and *R* (*Round ascospores*), in which known or suspected deletion alleles induce MSUD in their wild-type counterparts (*ban*^*+*^, *dip-1*^*+*^, *pk*^*+*^, and *r*^*+*^). The *Sad-1* and *Sad-2* suppressors also suppressed the barren phenotype of crosses heterozygous for chromosome segment duplications (i.e., *Dp* x *N*) (Kasbekar, 2013; Shiu *et al*., 2001; 2006). *Dp(EB4)* and *Dp(IBj5)* strains contain duplications of, respectively, 35-and 115-gene segments (Kasbekar 2013). Crosses of *Dp(EB4)* and *Dp(IBj5)* strains with the OR type wild strains were barren, with the Sad type were fertile, and with the Esm type were, respectively, fertile and barren (Ramakrishnan *et al*., 2011). Reduced MSUD in the related species *N. tetrasperma* also was attributed to asynapsis and silencing of the *sad-1* gene because of structural difference between the mating-type chromosomes (Jacobson *et al*., 2008). An alternative hypothesis (model 2) posits that natural populations harbor a wide variation in MSUD strength, and the OR strains represent the MSUD-conducive extreme. Model 1 predicts that crosses isogenic for a wild-isolated Sad type genetic background would show an OR-like MSUD phenotype. Here, we examined MSUD in tester-heterozygous crosses made in a novel isogenic background generated from the Sad type *N. crassa* wild strains Bichpuri-1 *a* and Spurger-3 *A*.

MSUD is also much less efficient in the *N. tetrasperma* strain 85 background (Jacobson *et al*., 2008; Ramakrishnan *et al*., 2011). Specifically, self-cross of an 85-derived [::*asm-1+*WT] dikaryotic strain produced < 76% MSUD-induced white-spored asci whereas comparable crosses in the OR background produced >99% white-spored asci (Ramakrishnan *et al*., 2011), thus the 85 background can be thought of as Sad-or Esm-type. We have now introgressed an ::*r* transgene from *N. crassa* OR into *N. tetrasperma* 85, and the resulting ::*r*^*Nt*^ testers were used to make ::*r*^*Nt*^ x 85 crosses. Additionally, by introgressing the *N. crassa T(EB4)* and *T(IBj5)* translocations into *N. tetrasperma* we had previously obtained *T(EB4)^Nt^, T(IBj5)^Nt^, Dp(EB4)^Nt^* and *Dp(IBj5)^Nt^* strains in which nominally only the rearrangement breakpoints were derived from *N. crassa* while the rest of the genome was derived from *N. tetrasperma* strain 85 (Giri *et al*., 2015). These strains enabled us to ask whether *Dp*-heterozygous crosses in *N. tetrasperma* are barren like their *N. crassa* counterparts.

In both *N. crassa* and *N. tetrasperma* the parental *mat A* and *mat a* haploid nuclei fuse during a sexual cross to produce a diploid zygote nucleus that via meiosis and a post-meiotic mitosis generates eight haploid progeny nuclei (4 *mat A* plus 4 *mat a*). In *N. crassa* these nuclei are partitioned into the eight initially uninucleate ascospores that form per ascus, whereas *N. tetrasperma* asci form four larger initially binucleate ascospores, each receiving a *mat A* and *mat a* pair (Raju, 1992; Raju and Perkins, 1994). Consequently, *N. crassa* mycelia are homokaryotic for mating type and require mycelium from another ascospore to complete the sexual cycle (heterothallic lifecycle), whereas dikaryotic [*mat A+mat a*] *N. tetrasperma* mycelia can undergo a self-cross (pseudohomothallic lifecycle). Dikaryotic mycelia can make some homokaryotic conidia (vegetative spores) by chance. Also, in *N. tetrasperma* ascogenesis, a pair of smaller homokaryotic ascospores can occasionally replace a dikaryotic ascospore to form a minor fraction of 5-8 spored asci. Mycelia from homokaryotic ascospores and conidia enable *N. tetrasperma* to out-cross with like mycelia of the opposite mating type. The *Eight spore* (*E*) mutant increases the replacement of dikaryotic ascospores by homokaryotic pairs, and *E* x *WT* crosses produce many 8-spored asci, although *E* x *E* crosses are infertile (Dodge 1939; Calhoun and Howe 1968). Does the ascus-dominant *E* mutant phenotype involve MSUD? An earlier attempt to generate *N. tetrasperma Sad-1* mutants did not yield an allele that could silence its *sad-1^+^* homologue (Bhat *et al*., 2004), therefore we introgressed a strong *N. crassa Sad-2* MSUD suppressor into *N. tetrasperma* 85 to enable us to compare *Sad-2* x *E* and *Sad-2^+^* x *E* crosses.

## Materials and methods

### Neurospora strains, culture methods, and crosses

Unless indicated otherwise, all Neurospora strains were obtained from the Fungal Genetics Stock Center (FGSC; McCluskey *et al*., 2010), Department of Plant Pathology, 4024 Throckmorton Plant Sciences Center, Kansas State University, Manhattan, KS 66506, USA. Neurospora was cultured essentially as described by Davis and De Serres (1970), using Metzenberg’s (2003) alternative recipe to make Vogel’s medium N. Crosses were made at 25 C by simultaneously inoculating mycelial plugs of the parental strains on synthetic cross medium supplemented with 1% sucrose and 2% agar.

*N. crassa*: The standard laboratory Oak Ridge strains 74-OR23-1VA (FGSC 2489) and 74-ORS-6a (FGSC 4200), referred to henceforth as OR *A* and OR *a*; and the wild-isolated strains Bichpuri-1 *a* (P750) and Spurger-3 *A* (FGSC 3201). The RLM 30-12 strain (genotype *Sad-2* (*RIP32*) *A*) containing a RIP-mutated *Sad-2* allele was a gift from Robert L. Metzenberg and is described by Shiu *et al.* (2006).

The OR-derived MSUD tester strains ISU 3118 (genotype: *rid his3*; *mus-52*Δ::*bar*; *VIIL::r^ef1^-hph A*) and ISU 3119 (genotype *rid his-3*; *mus-52*Δ::*bar*; *VIIL*::*r*^*ef3*^-*hph A*) bearing the transgenes ::*r1* and ::*r3* were a gift from Dr. Tom Hammond, Illinois State University, Normal, IL 61790, USA, and are described by Samarajeewa *et al.* (2014). The 5.5 kb long *r*^*+*^ gene (ncu02764, nucleotides 9,280,963 to 9,286,536) on chromosome 1 encodes a 3.3 kb ORF that shows robust MSUD that is suppressible only by the strong *Sad-1* and *Sad-2* suppressors, unlike other genes (*act, asm-1, mei-3*) in which MSUD can be suppressed by even the weaker MSUD suppressors (Hammond *et al*. 2013a). The testers obtained from Dr. Hammond are henceforth referred to as ::*r1*^*OR*^ and ::*r3*^*OR*^ to distinguish them from the ::*r1^B/S1^* and ::*r3^B/S1^* testers that we constructed in the B/S1 genetic background (described below). The insertion sites of the ::*r1^B/S1^*and ::*r3^B/S1^* transgenes are the same as those of ::*r1*^*OR*^ and ::*r3*^*OR*^.

Strains #1 (genotype ::*r1^OR^a*) and #6 (genotype ::*r1^OR^A*) were obtained as segregants from the cross ISU 3118 x OR *a*, and strains #1 and #10 (genotype ::*r3^OR^a*), and #11 (genotype::*r3^OR^A*) were obtained from ISU 3119 x OR *a*. Strains #4 (genotype ::*r1^B/S1^ a*) and #1 (genotype ::*r1^B/S1^ A*) were obtained from ::*r1^B/S1^ A* x B/S1 *a*, and strains #4 and #16 (genotype::*r3^B/S1^A*) and #15 and #24 (genotype ::*r3^B/S1^ a*) were obtained from ::*r3^B/S1^ A* x B/S1 *a* (described below). The results of crosses with the ::*r1*^*OR*^ and ::*r1^B/S1^* strains are presented in Table S2, and those with ::*r3*^*OR*^ and ::*r3^B/S1^* are presented in Table 1.

*N. tetrasperma*: The reference strains 85 *A* (FGSC 1270) and 85 *a* (FGSC 1271); the *E* mutants *lwn; al(102), E A* (FGSC 2783) and *lwn; al(102), E a* (FGSC 2784) (hereafter *E A* and *E a*). Previously, we had introgressed the *N. crassa T(EB4)* and *T(IBj5)* translocations into *N. tetrasperma* to construct the strains *T(EB4)^Nt^* and *T(IBj5)^Nt^*, and from the *T*^*Nt*^ x 85 crosses we obtained self-fertile dikaryotic [*T*^*Nt*^*+N*] and [*Dp*^*Nt*^*+Df*^*Nt*^] strains (Giri *et al*., 2015). From the dikaryons we obtained homokaryotic conidial derivatives of genotype *T(EB4)^Nt^ a, T(IBj5)^Nt^ a, Dp(EB4)^Nt^ A* and *Dp(IBj5)^Nt^ a* and used them to make the crosses whose results are presented in Table 3.

The *Sad-2^Nt^* and ::*r3*^*Nt*^ strains were made by introgressing the relevant *N. crassa* gene into the strain 85 genetic background (see below). They were used to measure MSUD efficiency in::*r3*^*Nt*^-heterozygous and ::*r3*^*Nt*^-homozygous crosses and in ::*r3*^*Nt*^ x *Sad-2^Nt^*. The superscript “*Nt*” in the strain designations (eg., ::*r3^Nt^ A, ::r3^Nt^ a, Sad-2^Nt^ A*, and *WT*^*Nt*^*A*) indicates that the strain has in its ancestry one or more self-fertile dikaryotic strain (i.e., *N. tetrasperma*-like).

*N. crassa/N. tetrasperma* hybrid strain: *C4,T4 a* (FGSC 1778; Metzenberg and Ahlgren, 1969) was used as a bridging strain to initiate the introgression of *N. crassa* genes into *N. tetrasperma*. Crosses of *C4,T4a* with opposite mating type strains of *N. crassa* and *N. tetrasperma* strains can produce small numbers of viable progeny.

### Ascospore harvests

Lid harvests: Ascospores were harvested by washing the lids of Petri plates in which the crosses were made with 1.5 ml water. The ascospore suspension was concentrated to 0.1-0.5 ml and an aliquot was counted in a hemocytometer. Plate harvests: 2 ml water was added to the agar surface and a rubber policeman or spatula was used to scrape off the perithecia, mycelia and ascospores. The suspension was transferred to a 2 ml tube, spun for 60 seconds in a table top centrifuge to pellet the ascospores and the supernatant with debris was removed. The ascospore pellet was re-suspended in 0.1-0.5 ml water and an aliquot was counted in a hemocytometer.

### Construction of MSUD testers in the B/S1 background

A cross was made between the wild-isolated strains Bichpuri-1 *a* and Spurger-3 *A* (B *a* x S *A*) and the f_1_ progeny were used to make four f_1_ *a* x f_1_ *A* sib-pair crosses that initiated four recombinant inbred lines. Within a line, in each generation sibling progeny of opposite mating types were crossed to produce the next generation (i.e., sibling f_1_ *a* x f_1_ *A* to produce the f_2_, then sibling f_2_ *a* x f_2_ *A* to produce the f_3_, etc). A recessive sterility-causing mutation became homozygous in the f_4_ and f_3_ generation of, respectively, lines 2 and 4 (C. Usha and D. P. Kasbekar, unpublished results), but we were able to reach the f_10_ generation in lines 1 and 3. A *mat A* and *mat a* strain pair of the f_10_ generation of line 1 (specifically, segregants #1 and #3) are referred to henceforth as B/S1 *A* and B/S1 *a*.

The *mus-51* gene in the B/S1 background was mutated by RIP (Selker, 1990). Strains mutant in *mus-51* are defective for non-homologous end joining, consequently, any transforming DNA can integrate only via homologous recombination (Ninomiya *et al*. 2004). A DNA construct bearing a 1683 bp *mus-51* segment (−205 to 1478 bp of the 2046 bp MUS-51 ORF) and the hygromycin-resistance (*hph*) cassette (Carroll *et al*. 1994) was transformed by electroporation into B/S1 *A* conidia, and ectopic integration of the transforming DNA created the transgenic *Dp(mus-51)* duplication. The *Dp(mus-51) A* primary transformant was crossed to B/S1 *a* and the progeny were used to perform a *Dp(mus-51)*-homozygous cross. Of 40 progeny examined from late harvested ascospores, one *mat a* progeny (#24) was found to contain several RIP-induced mutations in the endogenous *mus-51* gene, including in-frame stop codons (Genbank accession number KM025239), and it was crossed with the B/S1 *A* strain to segregate out the *Dp(mus-51)* transgene and obtain the B/S1 *mus-51 A* #8, B/S1 *mus-51 a* #3 strains in the progeny.

Transformation of B/S1 *mus-51A* conidia was used to make the *tester*^*B/S1*^ strains, namely, ::*r1^B/S1^* and ::*r3^B/S1^*, which are analogous to the OR-derived ::*r1*^*OR*^ and ::*r3*^*OR*^ testers of Samarajeewa *et al.* (2014). A 2.3 kb fragment from the 3’ end of the *r*^*+*^ gene (*r*^*ef*^) was joined to the *hph* cassette by double-joint PCR (Yu *et al*. 2004) to create a 4.1 kb *r*^*ef*^-*hph* fusion construct that also included flanking sequences for its precise insertion by homologous recombination into the same genomic sites as in the ::*r1*^*OR*^ and ::*r3*^*OR*^ testers. The DNA constructs were transformed by electroporation into the B/S1 *mus-51 A* strain, and transformants were selected on hygromycin-medium. The potentially heterokaryotic primary transformants were crossed with B/S1 *a* to segregate out the *mus-51* mutation and obtain a ::*r3^B/S1^ A* homokaryon from whose cross with B/S1 *a* we obtained the segregants #15 and #24 (of genotype ::*r3^B/S1^ a*) and #4 and #16 (of genotype ::*r3^B/S1^A*). In a like manner we obtained an ::*r1^B/S1^ A* homokaryon that was crossed with B/S1 *a* to obtain the segregant #4 (of genotype ::*r1^B/S1^ a*) and segregant #1 (of genotype ::*r1^B/S1^ A*). Between derivation of the B/S1 *A* and *a* strains and the ::*r*^*B/S1*^ *A* and *a* strains five additional backcrosses were done to B/S1, thus increasing the chance that a ::*r*^*B/S1*^ *A* x ::*r*^*B/S1*^ *a* cross is further reduced in heterozygosity compared to a B/S1 *A* x B/S1 *a* cross.

Strains B/S1 *A* #1 (FGSC 26446), B/S1 *a* #3 (FGSC 26445), B/S1 *mus-51 A* #8 (FGSC 26444), B/S1 *mus-51 a* #3 (FGSC 26443), VIIL ::*r1^B/S1^-hph A* #1 (FGSC 26442), VIIL ::*r1^B/S1^-hph a* #4 (FGSC 26441), VIIL ::*r3^B/S1^-hph A* #16 (FGSC 26440), and VIIL ::*r3^B/S1^-hph a* #15 (FGSC 26439) have been deposited in the FGSC, with the accession numbers indicated in parenthesis.

### Derivation of the ::*r3*^*Nt*^ and *WT*^*Nt*^*A* strains

Strain ISU 3119 (genotype *VIIL::r3 A*) was crossed with the *N. crassa/N. tetrasperma* hybrid bridging strain C4T4 *a* and an ::*r3^1C^A* progeny was back-crossed with C4T4 *a* to produce an ::*r3^2C^ a* progeny, that was crossed with 85*A*, and an ::*r3^185^a* progeny was back-crossed again with 85 *A*. (Superscript “1C” indicates progeny from the first cross with C4T4 *a*, “2C” indicates progeny from the second cross with C4T4 *a*, and “n85” indicates progeny from the nth backcrosses with strain 85.) From the cross ::*r3^185^a* x 85 *A* we obtained several self-fertile progeny bearing the ::*r3* transgene, viz., 1R, 2R, 6R, 8R, 10R, 13R, 15R, 17R, and 20R. Self-cross of these self-fertile strains produced mostly four-spored asci, including several with round ascospores (% round ascospores, respectively, 90, 5, 5, 45, 90, 1, 2, 55, and 1). The crosses whose results are summarized in Table 2 were made with self-sterile conidial derivatives (CD) obtained from these self-fertile strains. The *VIIL::r3^Nt^ a* strain used was CD #4 derived from 10R, and *VIIL::r3^Nt^ A* was CD #5 derived from 1R. The ::*r3^Nt^ a* strain was deposited in the FGSC with the accession number FGSC 26489.

The *WT*^*Nt*^*A* strain in Table 2 was CD #5 derived from 6R, and it contains the non-transgenic nucleus of the 6R dikaryon. It was used as an additional “wild type” control. CD #1 derived from 13R turned out to be a self-sterile [::*r3^Nt^A+WT^Nt^A*] dikaryon whose identification and possible provenance are described in the Results and Discussion sections. The oligonucleotide primers 5’ATTCAATATCTGGCTTATCGATACCGTCCACCTC and 5’ACAGCGAACGAAACCCCTGAAAC were used to PCR amplify a 1.5 kb segment from genomic DNA of *N. crassa* and *N. tetrasperma* strains containing the chromosome 7 transgene ::*r3*. Strains bearing a transgene-free chromosome 7 were detected by PCR amplification of a 2.5 kb amplicon with the primers 5’GTTGAGGGTCTTGAGGGCGAAG and either 5’CGAGGGCCGAGTCTGGTGGTTA (for OR-derived DNA) or 5’CGAGGGCCGTGTCTGGTGGTTG (for strain 85-derived DNA).

### Derivation of the *Sad-2^Nt^* strains

The primers 5’GtAGCCAAGCTCtATtATtGtACtGCCTat (lower case indicates RIP-altered bases) and 5’ATCGATGGGAGCCAAGTGTAT can amplify a 1 kb segment in PCR using genomic DNA of strains bearing the *Sad-2* (*RIP32*) allele as template, whereas primers 5’CTGTGCGAGAACTTGATGATCTTGGAGGCG and 5’GAGCTTTGGCGTAAATCAGTTGACG can amplify a 619 bp amplicon in PCR with genomic DNA of *N. crassa* and *N. tetrasperma* strains containing the wild type *sad-2^+^* allele.

From the cross of the *N. crassa* RLM 30-12 strain (genotype *Sad-2* (*RIP32*) *A*) with *C4,T4 a* we identified a *Sad-2^1C^ a* progeny by PCR, and from the cross *Sad-2^1C^ a* x 85*A* we obtained a *Sad-2^185^ a* segregant and used it to initiate a series of backcrosses with the 85*A* or 85 *a* strains. After six backcrosses we obtained a self-sterile [*Sad-2^685^ A+Sad-2^685^ a*] dikaryon that was productive in crosses with both 85 *A* and 85 *a* (Figure S1; bent arrows in the figure indicate identification of the *Sad-2* progeny by PCR). From the dikaryon’s cross with 85 *a* we obtained S12, a self-fertile dikaryon of presumed genotype [*Sad-2^785^a+sad-2^+^ A*]. From the self–cross of S12 we obtained the [*Sad-2+sad-2*^+^] dikaryons 1S12 and 3S12, and, from the self-cross of 1S12 we obtained the [*Sad-2+sad-2*^+^] self-fertile dikaryons 1(1S12), 3(1S12), 4(1S12), 9(1S12). The *Sad-2^Nt^A* strain used in Table 2 is a *mat A* conidial derivative from 3(1S12). To derive the *Sad-2^Nt^a* strain we obtained one *Sad-2 a* homokaryotic progeny (#20) from the cross of the self-sterile [*Sad-2^685^ A+Sad-2^685^ a*] dikaryon with 85 *A*. This cross yielded mostly self-fertile progeny, including one called S1, from which we obtained the *Sad-2^Nt^ a* strain as a conidial derivative, and it is deposited in the FGSC with accession number FGSC 26490.

### Whole genome sequence analysis of Neurospora strains

DNA samples were sequenced at Fasteris, Switzerland, on Illumina HiSeq 4500 platform with paired end reads and read length of 150 bp. Quality and statistics of the raw reads were analyzed using FastQC (Version 0.11.5). Illumina adapters from the paired reads were removed using trimmomatics (version 0.36) (Bolger *et al*., 2014). Preprocessed reads were aligned to the *N. crassa* OR74A reference genome (version NC12, release 38, ensembl) using bowtie2 (version 2.3.4) with end-to-end parameter (Langmead and Salzberg, 2012). All samples were observed to have more than 80% alignment rate and at least ∽ 8 million aligned reads (Table S1). Duplicated reads in the alignment file were removed using picardtools (version 2.17.4). Variant calling or genotyping of the eight samples (NC1 to NC8) was performed using Gatk HaplotypeCaller (version 4.0.0.0) (DePristo *et al*., 2011; Van der Auwera *et al*., 2013). We filtered out SNPs with QD (Quality by depth) < 20. Genotype comparison among the NC1 to NC8 samples were performed using custom R script (version 3.4.3). The whole genome sequence data for the NC1 to NC8 strains are available from the National Centre for Biotechnology Information Sequence Read Archive under the accession number SRP149022.

## Results

### Estimates of genome heterozygosity in the near-isogenic ::*r3^B/S1^* x B/S1 crosses

In the series of sib-pair crosses made to produce the B/S1 strains, heterozygosity was selected for in each generation only at the *mat* locus, and nowhere else (see Materials and methods). Therefore a B/S1 *A* x B/S1 *a* cross is expected to be heterozygous in only ∽0.1% (i.e. ∽43 kbp) of the *mat*-unlinked genome fraction, since the residual heterozygosity is halved in each successive sib-pair cross (100%, 50%, 25%, …, 0.4%, 0.2%, 0.1%). To verify this conclusion we generated a genome wide high resolution map of residual heterozygosity in the B/S1 *A* x B/S1 *a* cross using Illumina high throughput sequencing. The whole genome sequence information of B/S1 *A* and B/S1 *a* identified 8784 SNPs in the *in silico* B/S1 *A* x B/S1 *a* cross. The number of heterozygous SNPs between B/S1 *a* and B/S1 *A* excludes the *mat* locus in which the *mat A* and *mat a* idiomorph sequences are very dissimilar seemingly unrelated. Of these, 7961 (90.6%) were in a 568 kb region that included the *mat* locus, 91 were elsewhere on chromosome I, and the remaining 732 were distributed on the other chromosomes (chromosome 2-387; 3-49; 4-104; 5-113; 6-20; and 7-59). Since the Bichpuri-1 *a* and Spurger-3 *A* strain genome sequences are not available, we could not determine the percentage loss of heterozygosity or map the LOH boundaries. However, the SNP density in the *mat*-flanking region is 14 SNPs /kb (7961 SNPs/ 568 kb), whereas in the rest of genome it is 0.02 SNPs/ kb (823 SNPs/ 39894 kb). The ratio of SNP density in the *mat*-unlinked region to that in the *mat*-linked region is 0.14% (0.02/14), which agrees with the 0.1% estimate of residual heterozygosity made above.

As pointed out in the Materials and Methods, construction of the ::*r*^*B/S1*^ *A* and *a* strains entailed doing five additional backcrosses to the B/S1 background, thus giving further chances to reduce heterozygosity in the ::*r1^B/S1^ A* x ::*r1^B/S1^ a* cross relative to B/S1 *A* x B/S1 *a*. A series of PCR-based Bichpuri-1 versus Spurger-3 RFLP markers in the chromosome 1 *mat*-distal (d) and *mat*-proximal (p) segments were found and we determined their homozygosity or heterozygosity (respectively, designated with the subscripts i and j) in a ::*r*^*B/S1*^ *A* x ::*r*^*B/S1*^ *a* cross, until the d_i_-d_j_ and p_i_-p_j_ intervals became small enough to be sequenced from the ::*r3^B/S1^ A* and ::*r3^B/S1^ a* DNA. Sequence alignment (Figure 1) revealed that a 258,732 bp segment, bounded by SNPs at positions 1629710 and 1888441 with respect to the OR genome sequence, was heterozygous in the ::*r3^B/S1^a* x ::*r3^B/S1^A* and the B/S1*A* x ::*r3^B/S1^a* crosses, whereas a 568 kb region identified in Illumina sequences was heterozygous in the B/S1*a* x B/S1*A* and B/S1*a* x ::*r3^B/S1^A* crosses. Thus, the B/S1*A* x ::*r3^B/S1^a* and B/S1*a* x ::*r3^B/S1^A* crosses are heterozygous for, respectively, ∽302 kbp (0.7%) and ∽603 kbp (1.4%) of the genome. Heterozygosity in the B/S1 x ::*r1^B/S1^* crosses also is not expected to exceed 1.4% of the genome.

**Figure 1:**
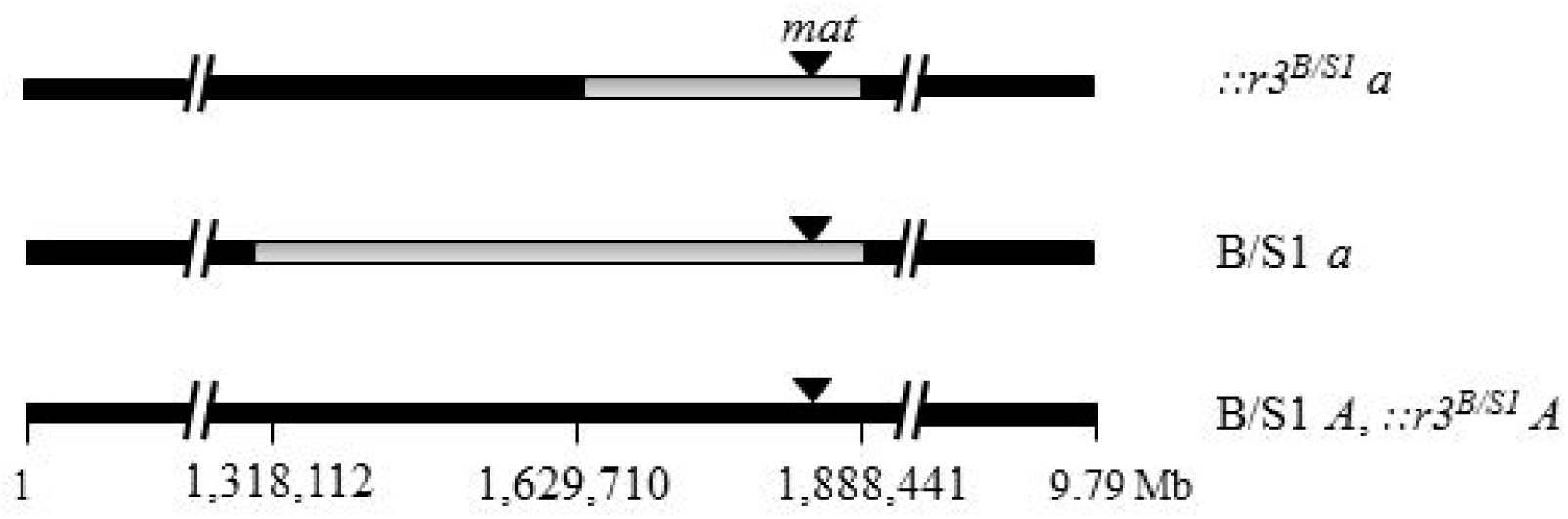
Heterozygosity in the *mat*-linked region of crosses in the B/S1 background. Chromosome 1 is 9.79 Mb long and its nucleotides are numbered as per the OR genome sequence (ID-CM002236). In the B/S1 *a* and::*r3^B/S1^ a* strains the *mat a* locus is derived from the Bichpuri-1 a strain. In the B/S1 *a* strain, sequences distal to nucleotide 1318112 and proximal to nucleotide 1888441 are identical to the B/S1 *A* sequence. Thus B/S1 *a* x B/S1 *A* and B/S1 *a* x ::*r3^B/S1^ A* crosses are heterozygous for a < 570329 bp segment between nucleotides 1318112 and 1888441. In the crosses ::*r3^B/S1^a* x ::*r3^B/S1^A* and B/S1 *A* x ::*r3^B/S1^ a* nucleotides at positions 1629710 and 1888441 represent heterozygous SNPs and the ∽258732 bp segment between them is heterozygous, whereas sequences distal to the former and proximal to the latter SNPs are homozygous. Sequence accession numbers are MG009253 and MG017489 (Spurger), MG009254 and MG017490 (B/S1 *A*), MG009255 and MG017491 (::*r3^B/S1^a*), and MG009256 and MG017492 (Bichpuri).

### MSUD efficiency differs in the *N. crassa* OR and B/S1 backgrounds

Model 1 (see Introduction) predicts that MSUD would be more efficient in B/S1 x ::*r*^*B/S1*^ crosses than in OR x ::*r*^*B/S1*^ or B/S1 x ::*r*^*OR*^, since the latter crosses are surely more heterozygous. The results summarized in Tables 1 and S2 lead us to reject this model. As can be seen in the Tables, four control crosses that did not contain any ::*r* transgene (OR *A* x OR *a*, B/S1 *A* x B/S1 *a*, OR *A* x B/S1 *a*, and OR *a* x B/S1 *A*) did not produce any round ascospores, although productivity of B/S1 *A* x B/S1 *a* was significantly lower than that of OR *A* x OR *a* and the two OR x B/S1 crosses. The ::*r3*^*OR*^ x OR crosses produced > 96 % round ascospores, while the homozygous ::*r3^OR^ a* x ::*r3^OR^ A* cross yielded 1 % round ascospores (Table 1), which confirmed the efficient MSUD characteristic of the OR genetic background. The two ::*r3^B/S1^* x ::*r3*^*OR*^ crosses produced < 4% round ascospores, showing that the MSUD machinery recognized the ::*r3^B/S1^* and ::*r3*^*OR*^ transgenes as alleles (paired). The ::*r3*^*OR*^ x B/S1 and ::*r3^B/S1^* x OR crosses produced 25-64 % round ascospores, which was consistent with the results of Ramakrishnan *et al*. (2011) showing MSUD is inefficient in crosses of the OR-derived MSUD testers with the Bichpuri-1 and Spurger-3 strains, from which the B/S1 strains are derived. The ::*r3^B/S1^* x B/S1 crosses produced < 24% round ascospores, which was inconsistent with model 1. Additionally, the homozygous ::*r3^B/S1^ a* x ::*r3^B/S1^ A* cross produced ∽ 1.5-fold more round ascospores than the ::*r3^B/S1^* x ::*r3*^*OR*^ crosses. Essentially similar results were obtained in crosses with the ::*r1*^*OR*^ and ::*r1^B/S1^* testers (Table S2). The results allow us to conclude that MSUD is less efficient in the B/S1 than the OR background, and there is possibly an increase in inappropriate silencing of paired genes in the B/S1 background.

The results of the ::*r*^*OR*^ x B/S1 and ::*r*^*B/S1*^ x OR crosses show that the “efficient MSUD” OR phenotype was recessive to the “inefficient MSUD” B/S1 phenotype (Tables 1 and S2). We crossed 49 f_1_ progeny from the B/S1 x OR cross with ::*r3*^*OR*^ strains of the opposite mating type and crosses with seven f_1_ progeny were found to produce > 90% round ascospores (Figure 2), suggesting that ∽1/8 of the f_1_ progeny had inherited the recessive OR-type phenotype. Thus, the OR phenotype might be determined by as few as three unlinked OR-derived loci.

**Figure 2:**
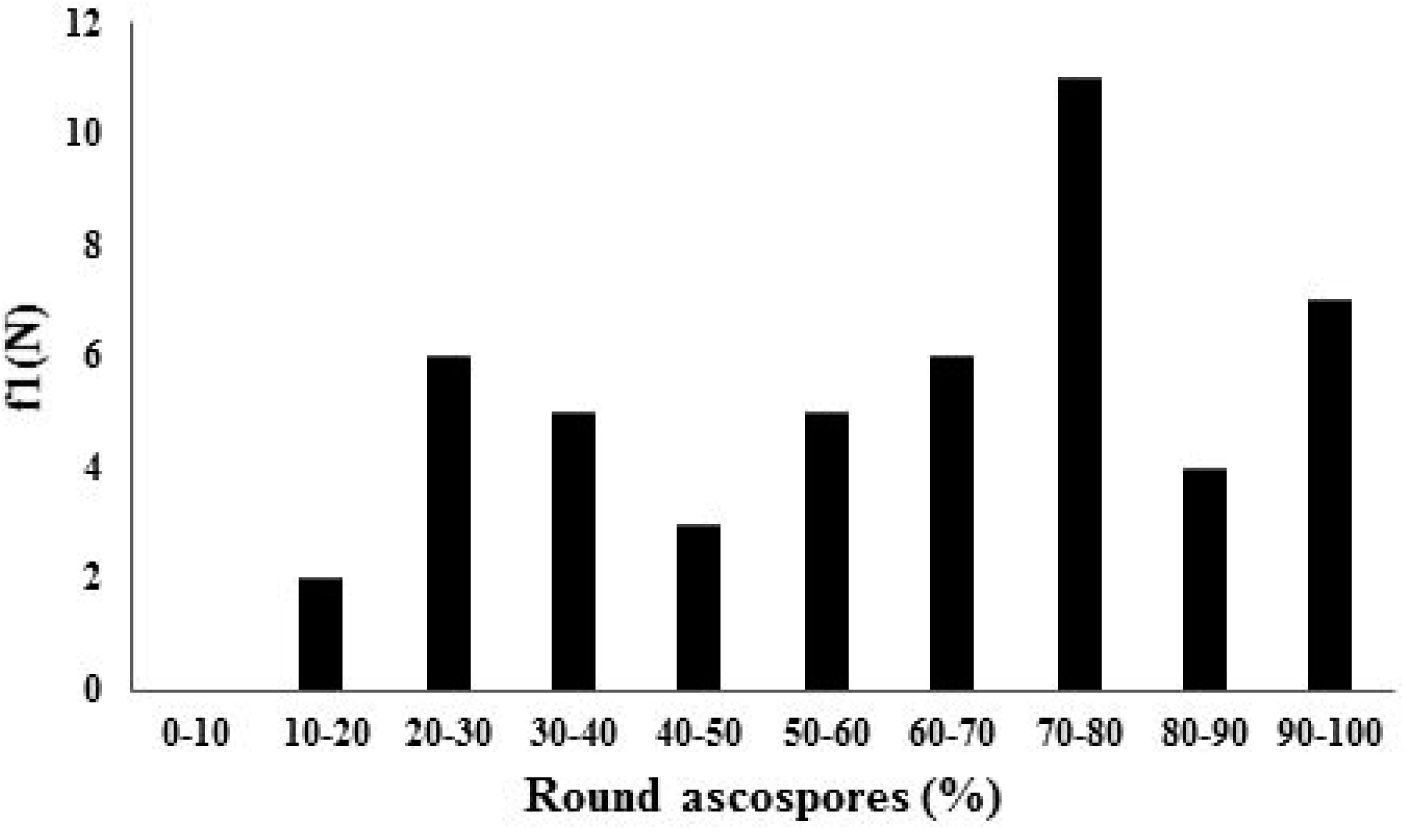
MSUD efficiency measured as the fraction of round ascospores produced in crosses of 48 f1 from B/S1 *A* x OR *a* with ::*r3*^*OR*^ strains of opposite mating type. The seven f1 progeny whose crosses produced >90 RSP are deemed to have inherited the efficient MSUD characteristic of the OR *a* parent. For six of the seven OR-type f1 progeny we examined the whole genome sequence to identify conserved OR-derived segments on chromosomes 1, 2 and 5.

To determine the loci responsible for the OR-type phenotype, we analyzed the whole genome sequence of six OR-type f_1_ progeny to identity conserved regions in their genomes with OR alleles. For this, we considered regions that showed the presence of OR allele in at least four of the six progeny, provided the other two did not show presence of the B/S1 allele. Based on these criteria, we observed that OR-derived alleles were fixed predominantly on chromosomes 1, 2 and 5 (Figure S2). These three chromosomal segments might harbor genes that contribute significantly to the OR-type phenotype.

### MSUD is inefficient in the *N. tetrasperma* 85 background

Table 2 summarizes the results of comparable crosses made in the *N. tetrasperma* strain 85 background. Again, control crosses not containing the ::*r3*^*Nt*^ transgene (85 *A* x 85 *a*, 85 *A* x *Sad-2^Nt^ a*, and *Sad-2^Nt^ A* x 85 *a*) did not produce any round ascospores, and the *Sad-2^Nt^ A* x *Sad-2^Nt^ a* cross was infertile like its *N. crassa* counterpart. The ::*r3*^*Nt*^ x 85 crosses produced ∽5-7% round ascospores, which was far less than the >96% seen in ::*r3*^*OR*^ x OR (Table 1). The ::*r3^Nt^a* x *WT*^*Nt*^*A* cross produced 21.5% round ascospores. The *WT*^*Nt*^*A* strain was a conidial derivative that carries the non-transgenic nucleus of the 6R dikaryon and it was used as an additional “wild-type” control. One ::*r3^Nt^A* x ::*r3^Nt^ a* cross and the cross *Sad-2^Nt^ A* x ::*r3^Nt^ a* yielded < 2.5% round ascospores, which suggested that MSUD does not occur in a transgene-homozygous cross, and that MSUD is suppressed by *Sad-2^Nt^*. However, a second putative ::*r3^Nt^A* x ::*r3^Nt^ a* cross, made using the strain CD #1 as the ::*r3^Nt^A* parent, yielded 11.3% round ascospores which was unexpectedly high for a presumed ::*r3*^*Nt*^–homozygous cross, and it was explored further as described below.

The crosses *Dp(EB4)^Nt^A* x 85 *a* and *Dp(IBj5)^Nt^a* x 85 *A* represent *Dp*-heterozygous crosses made in the *N. tetrasperma* 85 background. Significantly, neither showed an obvious barren phenotype, and they were only quantitatively less productive than the corresponding control crosses *T(EB4)^Nt^a* x 85 *A* and *T(IBj5)^Nt^a* x 85 *A* (Table 3). Finally, the *Sad-2^Nt^* x *E* and *E* x *Sad-2^+^* crosses showed no difference in ascus development (data not shown).

### A self-sterile dikaryon recovered from a self-fertile strain

The second putative ::*r3^Nt^A* x ::*r3^Nt^ a* cross that yielded an unexpectedly high fraction of round ascospores was made using the strain CD #1 as the ::*r3^Nt^A* parent. CD #1 was a self-sterile conidial derivative obtained from the self-fertile strain 13R (see Materials and Methods), and we considered the possibility that it was in fact a [(::*r3^Nt^A*)+(*+A*)] dikaryon. In which case its cross with the ::*r3^Nt^ a* strain would represent two crosses, one of genotype ::*r3^Nt^A* x ::*r3^Nt^ a*, and the other of genotype (*+A*) x ::*r3^Nt^ a*, and most of the round ascospores would come from the latter. PCR with CD #1 genomic DNA and primers specific for either a transgene-bearing or transgene-free chromosome 7 revealed that such indeed was the case. Below we discuss the possible provenance of a dikaryotic [(::*r3^Nt^A*)+(*+A*)] conidial derivative from a self-fertile strain.

### Round ascospores show impaired dispersal

Ordinarily ascospores are harvested from the lids of the cross plate. This is simple enough to do and one can obtain adequate ascospore numbers. However, to obtain the results in Tables 1, 2, and S2 we combined the ascospores harvested from the lids together with those harvested from the agar surface of the cross plates and obtained a “total” productivity count. When we harvested ascospores from only the lids of the cross plates we found a significant deficit of round ascospores, especially in crosses with the B/S1-derived strains (Tables S3 and S4). Many round as well as spindle-shaped ascospores were found adhered to the outside of perithecia, or deposited close by on the agar, in crosses with the B/S1-derived strains and in *N. tetrasperma*. Thus, scoring only the lid-harvested ascospores would underestimate MSUD strength. These results suggest that round ascospores suffer impaired dispersal relative to the spindle-shaped ascospores, and the former might also interfere with dispersal of the latter, possibly by clogging the perithecial exit pore (ostiole). However, this effect is not as evident in the OR background, and as far as we are aware such difference between round and spindle-shaped ascospores was not previously reported.

## Discussion

### Efficient MSUD is not necessarily the norm in Neurospora

The *N. crassa* B/S1 and *N. tetrasperma* 85 genetic backgrounds exhibited two MSUD anomalies relative to the *N. crassa* OR background: (1) an inefficient silencing of unpaired genes, and (2) an apparent increase in inappropriate silencing of paired genes. While in the OR background the MSUD response in crosses heterozygous or homozygous for an ::*r* transgene was all-or-none (i.e., respectively, > 97% vs. < 1 % round ascospores), the results were not as clear-cut in comparable crosses in B/S1 and 85, and the round ascospore fractions were, respectively, 24 % vs. 6 %, and 7 % vs. 1.4 %. In fact, MSUD efficiency in the B/S1 background was comparable to that in OR crosses deficient for either the *sad-6^+^* gene, which encodes a Rad54-like SNF2 helicase-related protein, or the *cbp-20^+^* and *cbp-80^+^* genes, that encode the cap-binding complex proteins associated with the 5’ cap of eukaryotic mRNA (Samarajeewa *et al*. 2014; Decker *et al*. 2017). Thus, efficient MSUD, so characteristic of the OR background, does not appear to be the norm in *N. crassa*. Indeed, the use of OR strains for genetic studies might have fortuitously facilitated (1) the discovery of MSUD and its semi-dominant *Sad* suppressors; (2) the discovery of ascus-dominant mutants such as *Ban, Dip-1, Pk*^D^, and *R*; and (3) *Dp* identification via the barrenness of *Dp*-heterozygous crosses. Efficient MSUD in the OR background even enabled Turner and Perkins (1982) to recover normal-sequence progeny from a cross between two chromosome 1 inversions (see Kasbekar, 2006).

Our preliminary mapping studies suggest that the OR versus B/S1 difference might be determined by as few as three loci on chromosomes 1, 2, and 5. It would be of interest to know whether the same loci underlie the difference between OR and other Esm-and Sad-type wild strains. The proteins encoded by these loci in the non-OR-type backgrounds might provide additional cues that calibrate the MSUD response. Their absence from the background OR would make the response atypically all-or-none. The round ascospore fraction from the ::*r3*^*Nt*^ x 85 crosses was 5-7%, and that from ::*r3^Nt^a* x *WT*^*Nt*^*A* was 21.5%. The *WT*^*Nt*^*A* strain served as an additional “wild type” control. It was derived from the 6R dikaryon, obtained during the introgression of the ::*r3*^*Nt*^ transgene from *N. crassa* OR into *N. tetrasperma* 85, and contains a transgene-free chromosome 7. It would be interesting to examine its genome for OR-derived segments.

### *Dp*-mediated RIP-suppression in natural populations

The results in Table 3 show that *T*-and *Dp*-heterozygous crosses are comparably productive in *N. tetrasperma* 85. This is in marked contrast to the generally severe barrenness of *Dp*-heterozygous crosses in OR. Since most studies on *Dp*s to date were done in the OR background it was generally assumed that *all Dp*-heterozygous crosses in Neurospora are barren, despite the fact that Sad-and Esm-type wild-isolates were used by us to increase progeny numbers in *Dp* x *N* crosses before the advent of the *Sad* suppressors (Fehmer *et al*. 2001). Now we show that inefficient MSUD in non-OR backgrounds can make *Dp*-heterozygous crosses non-barren. Large *Dp*s (> 300 kbp) can act as dominant suppressors of RIP, presumably by titrating out the RIP machinery (Bhat and Kasbekar, 2001; Bhat *et al*., 2003; Vyas *et al*., 2006; Perkins *et al*., 2007; Singh and Kasbekar, 2008; Singh *et al*., 2009). Given that *Dp*-heterozygous crosses are not always barren, *Dp*-mediated RIP suppression might be more significant than is generally appreciated. For example, it can explain the persistence of the retrotransposon *Tad* in the *N. crassa* Adiopodoumé strain while all other Neurospora strains examined contain only RIP-inactivated relics. Possibly, an ancestral *Dp* strain sheltered *Tad* from RIP until its copy number increased sufficiently to render further *Dp*-mediated titration of the RIP machinery superfluous, and the *Dp*–generating rearrangement might itself have come from crossover between *Tad* copies (Kasbekar, 1999; 2013).

### Does MSUD underlie the ascus-dominant *E* phenotype?

There was no difference in ascus development in the crosses *Sad-2^Nt^* x *E* and *E* x *Sad-2^+^,* therefore it would appear that MSUD does not underlie the ascus-dominant *E* phenotype. However, this pre-supposes that silencing of *sad-2*^+^ by the semi-dominant *Sad-2* mutation is as effective in the 85 background as in OR. Consistent with this supposition the ::*r3*^*Nt*^ x 85 and *Sad-2^Nt^ A* x ::*r3^Nt^ a* crosses produced, respectively, 5-7% and < 2% round ascospores. Admittedly, a more definitive answer might come from examining *E* in *N. tetrasperma* crosses homozygous for either *sad-4Δ* or *sad-5Δ*. The SAD-4 and SAD-5 proteins are required in *N. crassa* for MSUD but, unlike SAD-2, they are dispensable for ascus development (Hammond *et al*., 2013a). The fact that certain combinations of intercrosses between single mating-type conidial derivatives derived from diverse wild-collected *N. tetrasperma* heterokaryons produced mostly eight-spored asci when one derivative was used as the female (protoperithecial) parent and mostly four-spored asci when the derivatives were reversed in the reciprocal cross (Jacobson, 1995), shows that eight-spored asci can form in *N. tetrasperma* independently of MSUD.

### Origin of the CD #1 conidial derivative

CD #1 was a self-sterile conidial derivative obtained from the *N. tetrasperma* self-fertile strain 13R. Its genotype was initially thought to be ::*r3^Nt^A*, but it turned out to be a [(::*r3^Nt^A*)+(*+A*)] dikaryon. Therefore the 13R strain likely had the trikaryotic genotype [(::*r3^Nt^A*)+(*+A*)+(*+a*)] or [(::*r3^Nt^A*)+(*+A*)+(::*r3^Nt^ a*)]. Self-cross of the trikaryon would include MSUD-induced round-spored asci, and it would generate self-sterile [(::*r3^Nt^A*)+(*+A*)] conidial derivatives. A trikaryotic ascospore can be generated if the ::*r3*^*Nt*^ locus undergoes second-division segregation, and one or more nucleus from the post-meiotic mitosis undergoes an additional mitoses before ascospore partitioning. We have recently reported that crosses involving hybrid strains made by introgressing *N. crassa* translocations into *N. tetrasperma* can occasionally undergo additional mitoses between the post-meiotic mitosis and ascospore partitioning, and thus produce more than eight nuclei in the pre-partition ascus (Kasbekar and Rekha, 2017). If more than eight nuclei are packaged into four ascospores then some will receive more than two nuclei. The recovery of the CD #1 strain cautioned us to re-confirm the genotype of all strains designated as homokaryotic in this work.

## Acknowledgements

We thank Ms. S. Rekha for assistance with some of the genetic crosses, and Ms. Parijat for processing the DNA samples for Illumina whole genome sequencing. DAG was supported by a CSIR-UGC Junior Research Fellowship. AVP was supported by a University Grants Commission fellowship (http://www.ugc.ac.in). KTN was supported by a Wellcome Trust-DBT India Alliance Intermediate Fellowship (IA/I/11/2500268) http://www.wellcomedbt.org and by Indian Institute of Science Education and Research Thiruvananthapuram intramural funds (http://www.iisertvm.ac.in). DPK is an INSA Senior Scientist in the Centre for DNA Fingerprinting and Diagnostics (CDFD), and Honorary Visiting Scientist in the Centre for Cellular and Molecular Biology (CCMB). A part of this work was supported by a grant from the SERB, Department of Science and Technology, Government of India, and by CDFD Core Funds to DPK during his tenure as Haldane Chair at CDFD. DPK thanks the CCMB for its hospitality.

